# Structural refinement of the auditory brainstem neurons in baboons during perinatal development

**DOI:** 10.1101/2020.12.23.424228

**Authors:** Eun-Jung Kim, Kaila Nip, Cynthia Blanco, Jun Hee Kim

**Author notes:** Corresponding author: Jun Hee Kim, Ph.D.

## Abstract

Children born prematurely suffer from learning disabilities and exhibit reading, speech, and cognitive difficulties, which are associated with an auditory processing disorder. However, it is unknown whether gestational age at delivery and the unnatural auditory environment in neonatal intensive care units (NICU) collectively affect proper auditory development and neuronal circuitry in premature newborns. We morphologically characterized fetal development of the medial superior olivary nucleus (MSO), an area important for binaural hearing and sound localization, in the auditory brainstem of baboon neonates at different gestational ages. Axonal and synaptic structures and the tonotopic differentiation of ion channels in the MSO underwent profound refinements after hearing onset in the uterus. In preterm baboon neonates, these developmental refinements of the MSO were significantly altered by limited maternal sound inputs from the isolated and unnatural environment in the NICU. Thus, the maternal environment, including auditory stimuli in uterus, is essential for auditory nervous system development during the last trimester of pregnancy and critically affects the anatomic and functional formation of synapses and neural circuitry in the preterm newborn brain.

## Introduction

Prematurity is one of the leading causes of perinatal mortality and long-term disability. Premature infants are at high risk for motor, sensory, cognitive, and behavioral disabilities ^1, 2^. In particular, neurosensory impairments such as deafness and visual deficits occur in 5-15% of preterm newborns ^3–5^. Auditory impairments caused by premature birth can profoundly affect reading and speaking ability and the development of language, all of which are associated with auditory processing disorder due to the inefficiency or incapacity of the auditory system to process acoustic information ^6, 7^. Preterm birth significantly increases the risk of neurodevelopmental disorders, such as autism spectrum disorder, that are characterized by some degree of auditory dysfunction including deafness, hyperacusis, difficulty listening in the presence of background noise, and problems in orienting and encoding speech sounds ^8, 9^. Notably, binaural hearing plays a key role in the development of speech and language perception because the normal development of the brain’s binaural circuitry requires the proper processing of auditory information ^10, 11^. For instance, a clinical study showed a delay in sound localization at 12 months was associated with deficits in the physiologic mechanism of temporal processing at an auditory processing evaluation done between 4 and 7 years of age ^12^. Thus, binaural hearing, sound localization, and temporal processing require more extensive central processing, the development of which is influenced by auditory experiences in different species from rodents to primates ^6, 13^.

The medial superior olivary nucleus (MSO) is an auditory brainstem nucleus and an important binaural site in the auditory pathway, where auditory inputs from the two ears converge and calculate the interaural time differences for computing sound location ^14^. The MSO is a prominent nucleus found in most small mammalian species. The biophysical, morphologic, synaptic, and biochemical properties of MSO neurons have been well described in gerbils, guinea pigs, mice, rats, and bats during their postnatal development ^15–17^. However, there are drastic differences in the developmental time-course and initiation factors of myelination and synaptic innervation between primates and rodents ^18–20^. Unlike mice and rats, human fetuses already have substantial capacity for auditory learning and memory ^6, 12, 19^, as they start to hear and undergo major developmental changes in the auditory system during the third trimester of pregnancy in the uterus ^19^. Understanding the neurophysiological effects of auditory experience on the MSO in the developing brain of non-human primate is essential to comprehend how experience-dependent plasticity drives the development of auditory processing in human.

Here, we investigated morphologic profiles of MSO neurons, including axonal structure, synapse, and tonotopic organization of ion channels throughout fetal development, specifically before and after hearing onset, in baboon neonates delivered at different gestational ages. Baboon fetuses begin to hear in the uterus at ~70% of gestational age (GA) and display five distinguishable waves of auditory brainstem responses (ABRs) at ~80% of GA ^21^, similar to humans. Baboons preferentially respond to lower-frequency sounds (<8 kHz, threshold: ~40 dB), as do human newborns. Studying the baboon brain thus allows direct comparisons with the human brain because the connectivity, size, and functional areas (reflected in motor and behavioral capacities) are highly similar to those in humans. Premature baboons delivered at 126 days (d) GA (70% of GA, term = 180 days GA), shares a similar neonatal course with human preterm infants delivered at 28 weeks of GA (extremely preterm), with common complications relevant to prematurity such as incomplete lung development and metabolism ^22^.

In addition, we studied how the altered auditory environment immediately after hearing onset influences the morphologic features of the MSO during early development. Preterm newborns experience a drastic change in going from the dark and muffled environment of the uterus to the unnatural environment of the NICU. During NICU stays, gestational age, medications, procedures, the unnatural auditory environment, and other stimuli can cause significant stress associated with adverse auditory development in preterm newborns ^18^. A difference of auditory environment between the maternal womb and the NICU, for instance attenuated maternal sound (e.g. maternal heart-beat sound) and increased unnecessary noise produced by ventilators, infusion pumps, or alarms, can cause potential risk for compromised neuroplasticity of the auditory nervous system in preterm newborns. Thus, we studied how altered auditory environment in an isolated environment in the NICU impacts the structural development and refinement of the MSO at the cellular level in developing baboon brains. These studies provide the first systematic description of MSO development at the cellular and synaptic level in non-human primates and reveals how premature birth alters the normal development of the MSO.

## Results

### Morphologic profiles of the MSO in baboon neonates during fetal development

Similar to the superior olivary complex (SOC) of humans ^23, 24^, the SOC from the baboon neonate brainstem delivered at 140d GA after hearing onset consists of the medial nucleus of the trapezoid body (MNTB), the MSO, the lateral superior olivary nucleus (LSO), the ventral nucleus of the trapezoid body (VNTB), and the lateral nucleus of the trapezoid body (LNTB). Baboon auditory brainstems labeled with MAP2 show prominent MNTB, MSO, and LSO organized in the medial to lateral directions in consecutive order. The MNTB was the most medial nucleus, with a round shape located along the auditory fibers that cross the midline of the brainstem ^25^. The MSO, a dorsoventrally orientated column of neurons, was located laterally to the MNTB. MAP2 staining showed MSO neurons with bipolar-like dendrites extending toward both the medial MNTB and lateral LSO. In the perinatal stage at 140d GA, individual neurons from those prominent nuclei (MNTB, MSO, and LSO) already had a typical morphology (e.g. bipolar structure of MSO neurons or multiple dendrites from LSO) and physiologic properties such as firing patterns (**Figure 1**).

**Figure 1.**
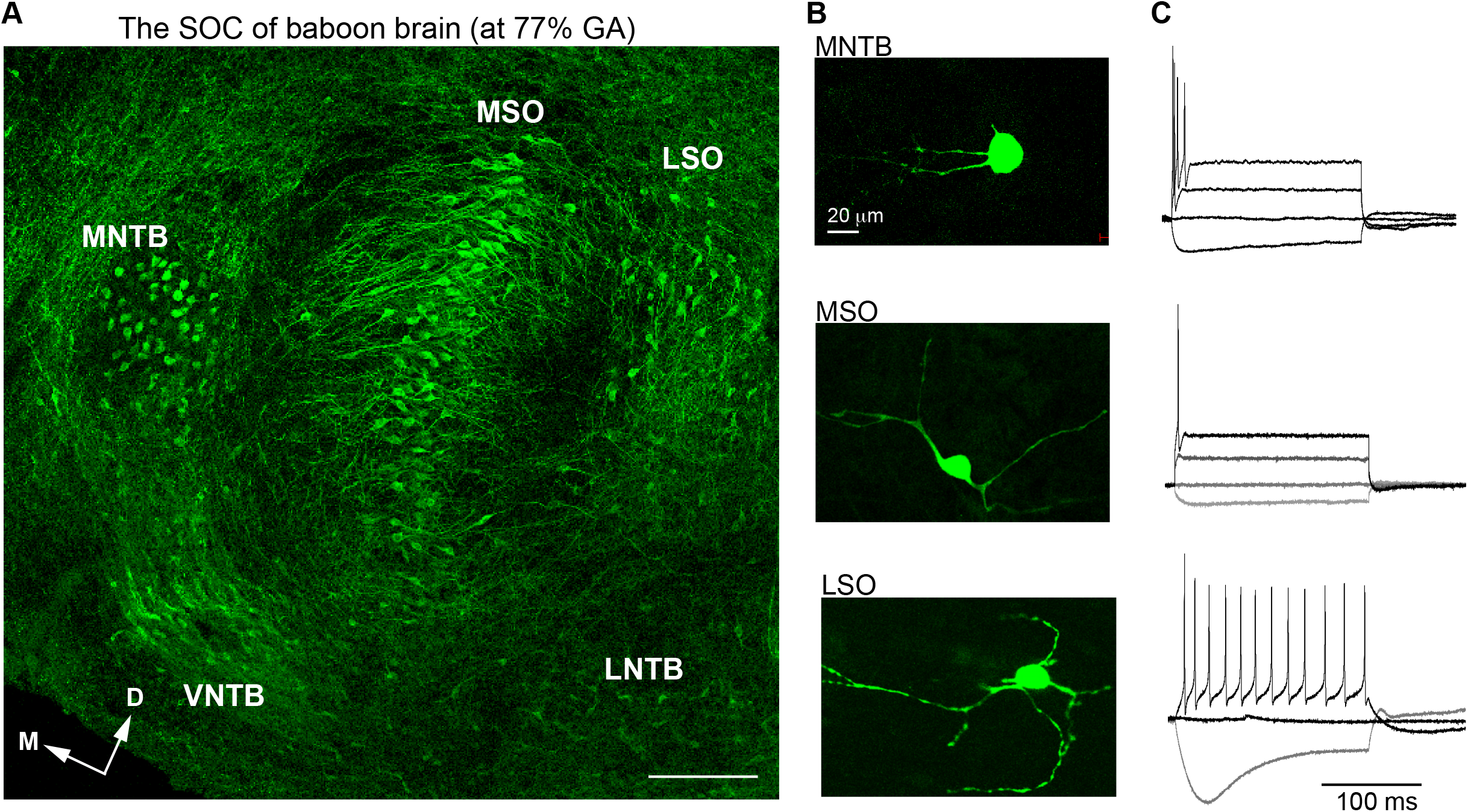
The main nuclei of the baboon SOC. **A.** The SOC in the auditory brainstem from a baboon neonate delivered at 77% GA, immunolabeled with MAP2 (green). From the midline of the brainstem, the MNTB, MSO, and LSO were arranged in a lateral direction. Scale = 200 μm. **B.** Individual cells from MNTB, MSO, and LSO nuclei were filled with Alexa dye during wholecell recordings. **C.** Action potentials from the same cells in B were evoked by step-like current injections (from −50 pA to 100 pA; top and middle, from −50 pA to 50 pA; bottom).

To determine the effects of auditory experience in the uterus on MSO development after hearing onset, we compared morphologic changes of the MSO in baboon neonates delivered at different gestational ages – extremely preterm, moderate preterm, and full-term birth; these corresponded to 67%, 77%, and 100% of 180-185 gestational days (full-term neonate, **Figure 2A**). Overall, the MSO became longer and narrower, transforming the outer line of MSO nuclei from an oval to a column shape with ventral-dorsal orientation during fetal development. In extremely preterm baboons (hereafter E-Pre) before hearing onset, the MSO was an oval shape (306,331 ± 22,794 μm^2^, n=7 slices from 5 baboons) and aligned along the ventral-dorsal axis.

**Figure 2.**
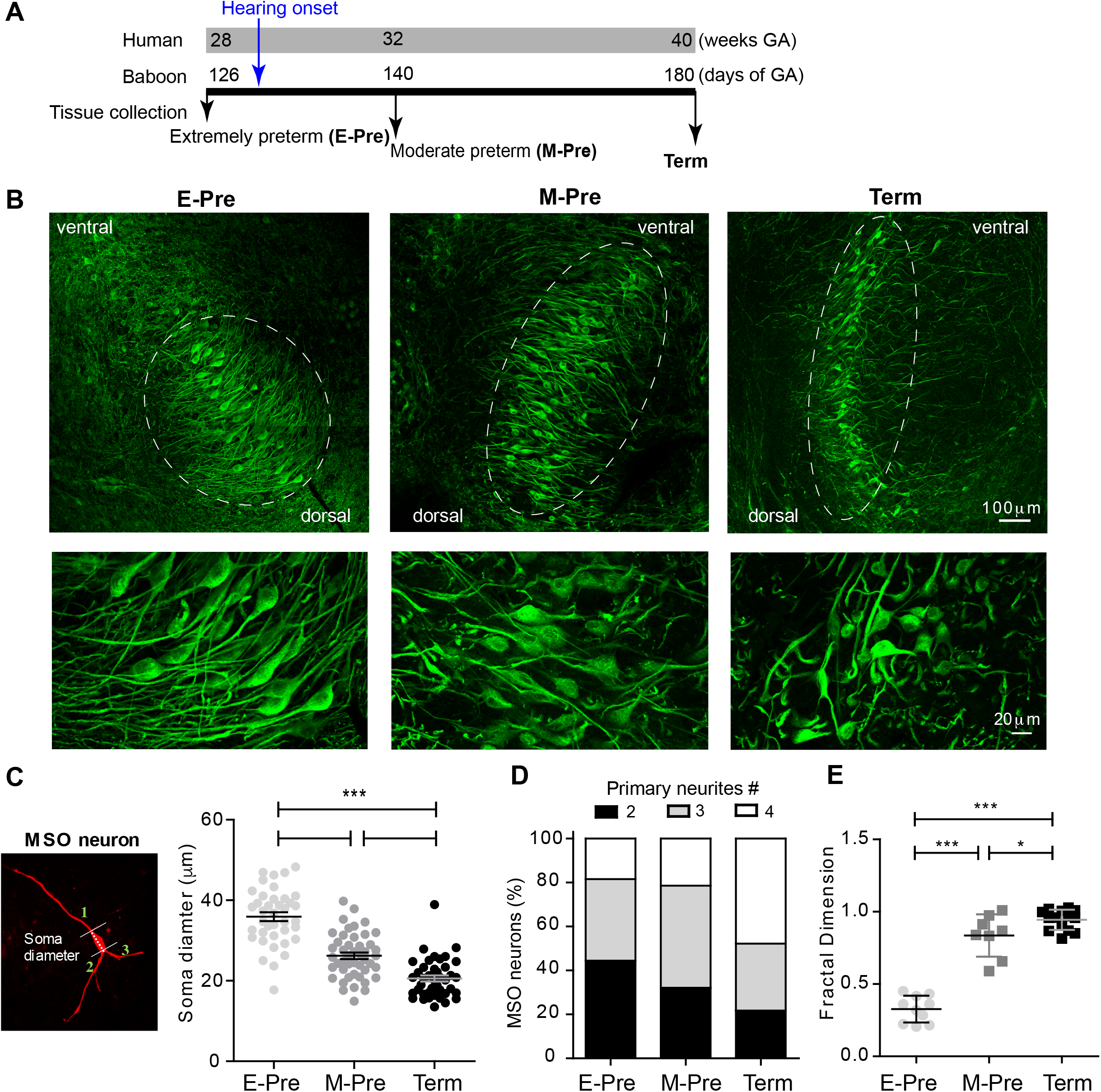
Morphologic changes in MSO neurons during gestational development of baboons in the uterus. **A.** Timeline for the last trimester of human (weeks) and baboon (days of gestational age, GA) gestation periods. Hearing onset is around 28 weeks of GA for human fetuses and after 126 days GA for baboons (blue arrow). Auditory brainstem tissues were collected from baboon neonates delivered at 126d GA (comparable to extremely preterm, E-Pre) and 140d GA (moderate preterm, M-Pre) via cesarean section, and full-term (Term) through natural delivery. Black arrows indicate the tissue collection time. **B.** MSO neurons were immunostained with MAP2 in the auditory brainstem from E-Pre, M-Pre, and Term neonates. (Top) Circles indicate the MSO area. Scale = 100 μm. (Bottom) Magnified image of MSO neurons. Scale = 20 μm. **C.** A single MSO neuron from a M-Pre neonate filled by Alexa dye during whole-cell recording. Broken line indicates the soma diameter and numbers indicate primary dendrites extending from the soma counted for quantitative analysis. Right: Somatic diameters from MSO neurons at different GA. **D.** Proportions of primary dendrites in MSO neurons from the soma at different GA. **E.** The fractal dimension, a measure of dendritic complexity, increased throughout the gestational ages in E-Pre, M-Pre, and Term neonates. Data were shown as mean ± s.e.m. *>0.05, **>0.01, and ***>0.001. One-way ANOVA and post-hoc Tukey’s multiple comparison test.

Most MSO neurons labeled with MAP2 exhibited a bipolar structure of dendrites that protruded toward both the medial MNTB and lateral LSO. But after hearing onset in moderate preterm baboons (hereafter M-Pre), the outer nuclei shape of the MSO highlighted by MAP2+ neurons became more condensed and narrower (283,045 ± 29,823 μm^2^, n=7 slices from 5 M-Pre baboons vs 306,331 ± 22,794 μm^2^, n=7 slices from 5 E-Pre baboons). In full-term baboons, the MSO was much thinner and longer than in preterm baboons (345,652 ± 21,317 μm^2^, n=11 slices from 3 baboons). MSO area per tissue was not significantly different in the gestational development groups, but the MSO neuron alignment along the dorsal-ventral axis was significantly longer in full-term baboons (hereafter Term, 1,074 ± 57.2 μm, n=8) compared to Epre (765 ± 62.4 μm, n=11, p=0.0025, Mann-Whitney test). The long axes of MSO neurons were arranged in a geometrically precise orientation with a dorsal to ventral direction (**Figure 2B**). These results reveal a developmental change in the shape of the MSO during the last trimester of pregnancy in baboon brains.

Next, we analyzed morphologic parameters of individual MSO neurons from preterm and full-term baboon neonates. The soma diameters of MSO neurons were 35.93 ± 0.101 μm (n=39) in E-Pre, 26.22 ± 0.819 μm (n=46) in M-Pre, and 20.57 ± 0.758 μm (n=41) in Term neonates (p<0.0001 by one-way ANOVA followed by Tukey post-hoc test) (**Figure 2C**). In E-Pre neonates, ~50% of MSO neurons had two primary dendrites, showing a clear bipolar structure. In M-Pre neonates, ~70% of MSO neurons had more than three primary dendrites extending from the soma. In Term neonates, ~80% of MSO neurons had multiple primary dendrites (**Figure 2D**). MSO neurons analyzed using theoretical fractal dimension analysis ^26^ were significantly more complex in M-Pre baboon neonates (0.83± 0.051, n=8) after hearing onset and further increased in Term neonates (0.94 ± 0.0185, n=14) compared to E-Pre baboons before hearing onset (0.32 ± 0.029, n=10, **Figure 2E**). Taken together these observations show that the MSO underwent significant morphologic changes in the outer shape of the nucleus, soma size, and the complexity of dendritic arbors in neurons after hearing onset in the uterus.

### Excitatory and inhibitory inputs on MSO neurons after hearing onset during fetal development in the uterus

Morphologic development of MSO neurons, including dendritic structure, may be regulated by synaptic inputs and integration ^27, 28^. We previously demonstrated that spatial refinement of synaptic inputs on the dendritic and somatic surfaces of the MSO neuron from baboon neonates occur after hearing onset ^25^. In the MSO of extremely preterm baboons, the vesicular GABA transporter (VGAT), which is a marker for inhibitory terminals, was visible on both cell bodies and dendrites; in term baboons, the MSO VGAT and vesicular glutamate transporter 1 (VGluT1, a marker for excitatory nerve terminals) puncta were concentrated on the soma and dendrites of MSO neurons, respectively (**Figure 3A, B**). Immunostaining of VGluT1 and VGAT showed that individual VGluT1 puncta area significantly increased in M-Pre MSO after hearing onset compared to Term neonates (1.38 ± 0.041 μm in E-Pre, 2.69 ± 0.142μm in M-Pre, 4.05 ± 0.084 μm^2^ in Term, p<0.0001 by one-way ANOVA followed by Tukey post-hoc test, **Figure 3C**). VGAT puncta area also increased (1.03 ± 0.034 μm^2^ in E-Pre, 2.14 ± 0.152 μm^2^ in M-Pre, and 3.02 ± 0.125 μm^2^ in Term, p<0.0001 by one-way ANOVA followed by Tukey post-hoc test, **Figure 3D**). However, the number of VGluT1 and VGAT puncta significantly decreased after hearing onset and remained this reduction throughout Term neonates. The number of VGluT1 puncta within a defined area (μm^2^) was 30.45 ± 4.765 (n=193) in E-Pre, 15.97 ± 3.851(n=159) in M-Pre, and 16.50 ± 3.267 (n=425) in Term (E-Pre vs M-Pre, p=0.0479 and M-Pre vs Term, p=0.9189, unpaired t-test). The number of VGAT puncta (number/ μm^2^) was 43.13 ± 6.936 (n=196) in E-Pre, 12.25 ± 2.398 (n=145) M-Pre, and 11.81 ± 0.988 (n=220) in Term neonates (E-Pre vs M-Pre, p=0.0024 and M-Pre vs Term, p=0.8443, unpaired t-test, **Figure 3E**). The increase in the size of VGluT1 and VGAT puncta could be due to changes in total vesicle number within individual boutons or distribution of synaptic vesicles around the active zone.

**Figure 3.**
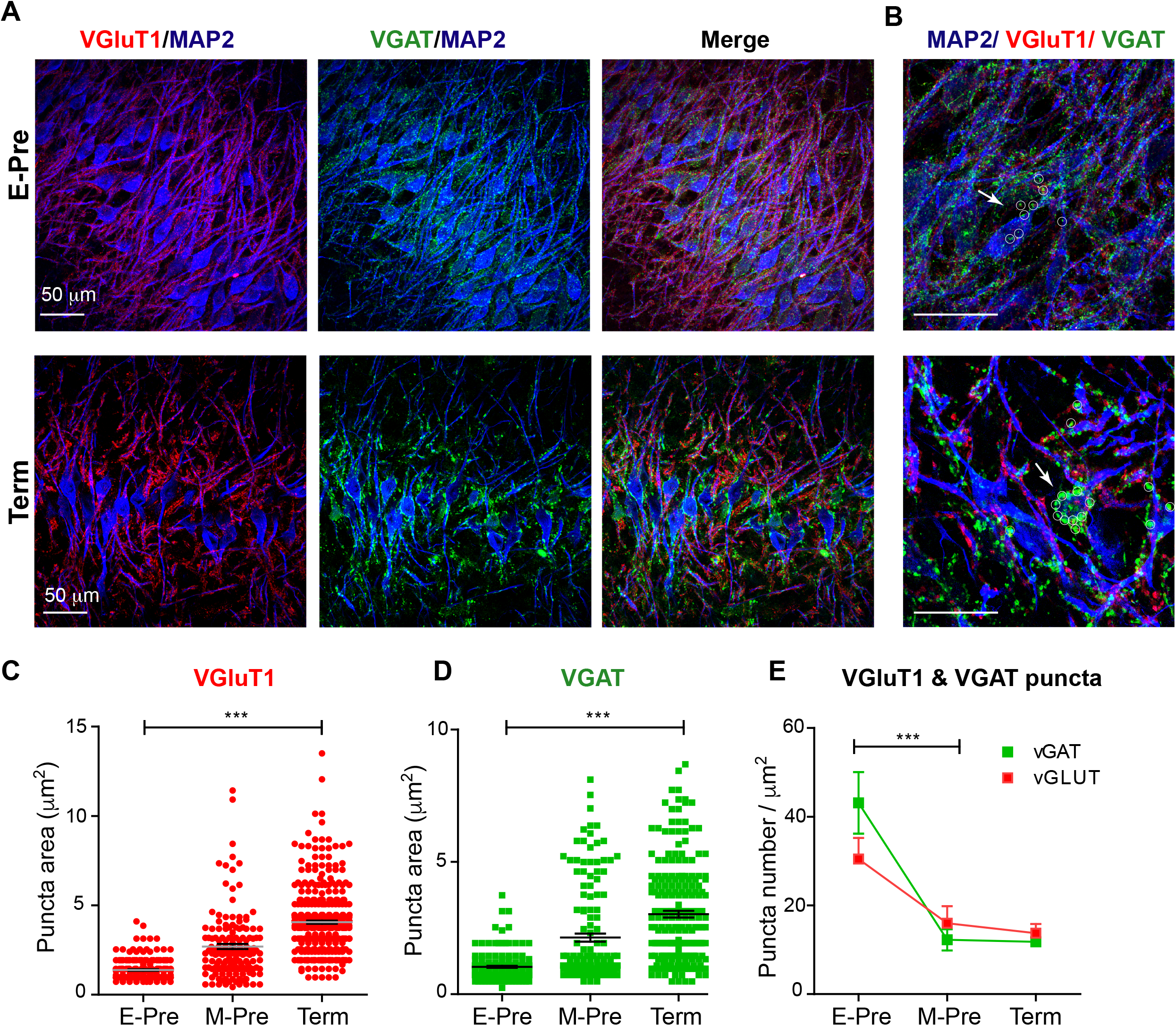
Synaptic inputs on MSO neurons during gestational development of baboon neonates. **A.** MSO nuclei were immunostained with MAP2 (blue), VGluT1 (red), and VGAT (green) in the auditory brainstem at E-Pre (top) and Term (bottom). Scale = 50 μm. **B**. Magnified image of MSO neurons. White arrows and small circles indicate VGAT signals (green) on MSO somata. Scale = 50 μm. **C-D.** Changes in the puncta area for VGluT1 (C) and VGAT (D) from E-Pre, M-Pre, and Term neonates. **E.** Numbers of VGluT1 and VGAT puncta per μm^2^ in the MSO.

Ultrastructural analysis of synapses in the MSO using transmission electron microscopy (TEM) showed no significant differences in the average active zone length (377.1 ± 17.68 nm, n=37 for E-Pre, 399.4 ± 18.29 nm, n=42 for M-Pre, and 436 ± 27.0 nm, n=20 for Term, p=0.175, one-way ANOVA and post-hoc Tukey’s multiple comparison, **Figure 4A, B**). However, the vesicles around the active zone (within 200 nm from the plasma membrane) and docked vesicles increased in Term neonates. Docked vesicles (within 10 nm) were 1.47 ± 0.24, n=36 for E-Pre, 1.98 ± 0.25, n=42 for M-Pre, and 3.0 ± 0.37, n=18 for Term (p=0.0052, one-way ANOVA and post-hoc Tukey’s multiple comparison, **Figure 4C**). The numbers of vesicles located within 100 nm (6.29 ± 0.68, n=37 for E-Pre, 6.33 ± 0.48, n=42 for M-Pre, or 10.5 ± 1.10, n=18 for Term) and 200 nm (11.58 ± 1.22, n=36 for E-Pre, 12.95 ± 0.88, n=42 for M-Pre, or 19.0 ± 1.83, n=18 for Term) were also increased throughout the gestational development (one-way ANOVA and Tukey’s multiple comparison, **Figure 4C**). In particular, the number of docked vesicles per the defined active zone (100 nm length, 0.34 ± 0.056, n=41 for E-Pre vs 0.82 ± 0.058, n=30 for Term, p<0.001, unpaired t-test) and total vesicle numbers (29.1 ± 3.41, n=34 for E-Pre vs 60.3 ± 5.92, n=18 for Term, p<0.001, unpaired t-test) were significantly increased in Term neonates compared with E-Pre neonates (**Figure 4D**). These results indicate that the distribution and strength of excitatory and inhibitory synaptic inputs to the MSO are dynamically refined by auditory inputs during the period of fetal development occurring after hearing onset in the uterus.

**Figure 4.**
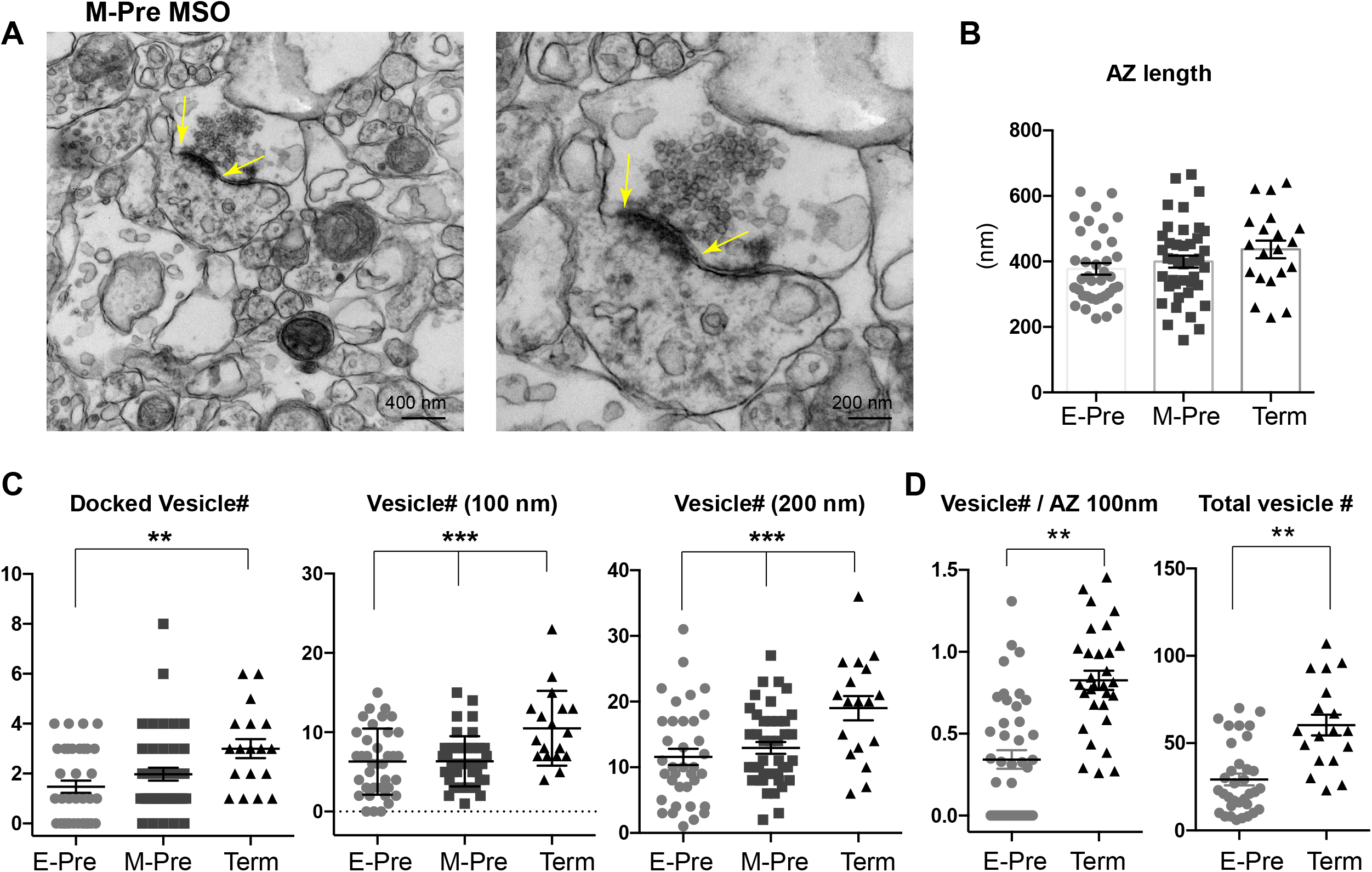
Presynaptic terminal undergoes structural refinement during gestational development. **A.** EM images of MSO synapse from M-Pre neonates. Yellow arrows indicate the active zone (AZ) at the presynaptic bouton. Scales= 400 and 200 nm. **B.** AZ length at the presynaptic boutons in the MSO from E-Pre, M-Pre, and Term neonate. **C.** The numbers of docked vesicles, which are located within 10 nm from the AZ, and numbers of vesicles located within 100 and 200 nm around the AZ in the presynaptic terminal. Data were shown as mean ± s.e.m. **>0.01 and ***>0.001, one-way ANOVA test. **D.** Numbers of docked vesicles in the defined AZ (100 nm) and numbers of total vesicles in individual boutons. Data were shown as mean ± s.e.m. **>0.01, unpaired t-test.

### Developmental refinement of K^+^ channel expression in the MSO neurons in the uterus

To determine whether developmental changes in synaptic and dendritic structures are accompanied with ion channel expression of MSO neurons, we examined K^+^ channel expression in MSO neurons. MSO neurons in gerbils, rats, and mice exhibit low- and high-K_v_ channels, which critically contribute to action potential firing in MSO neurons ^17, 27, 29, 30^. We examined expression of low-voltage-activated K_v_1.2 and high-voltage-activated K_v_3.1 channel in MSO neurons during gestational development. In M-Pre neonates, K_v_3.1 channel was expressed in the somatic and primary dendritic membranes of MSO neurons, whereas K_v_1.2 channel expression was restricted to the proximal axons of MSO neurons (**Figure 5A**). K_v_3.1 channel was broadly expressed in the somatic and primary dendritic membranes of MSO neurons in E-Pre throughout Term baboons, whereas K_v_1.2 channel expression underwent a developmental refinement. Notably, the length of the observed K_v_1.2 expression in the proximal axon became shorter specifically in ventral MSO neurons during fetal development (29.3 ± 1.13 μm, n=22 for E-Pre, 23.9 ± 1.12 μm, n=21 for M-Pre, and 19.6 ± 0.99 μm, n=17 for Term, p<0.0001, one-way ANOVA). There was no significant difference in K_v_1.2 expression in dorsal MSO neurons (**Figure 5B**). Therefore, after hearing onset in M-Pre and Term baboons, K_v_1.2 expression was significantly shorter in ventral MSO neurons rather than dorsal MSO neurons, indicating a tonotopic differentiation of K_v_1.2 channel expression in MSO neurons. In addition, the observed location of the K_v_1.2 channel expression became proximate to the soma of MSO neuron and more superimposed with the immunolabeled with ankyrinG (AnkG), representing the axon initial segment (AIS), in ventral MSO neurons during perinatal development (**Figure 5C-D**). In E-Pre MSO, K_v_1.2 expression was partially superimposed on AnkG expression and located at more distal axons. In MSO of Term baboons, the length of K_v_1.2 expression became shorter and almost co-localized with AnkG.

**Figure 5.**
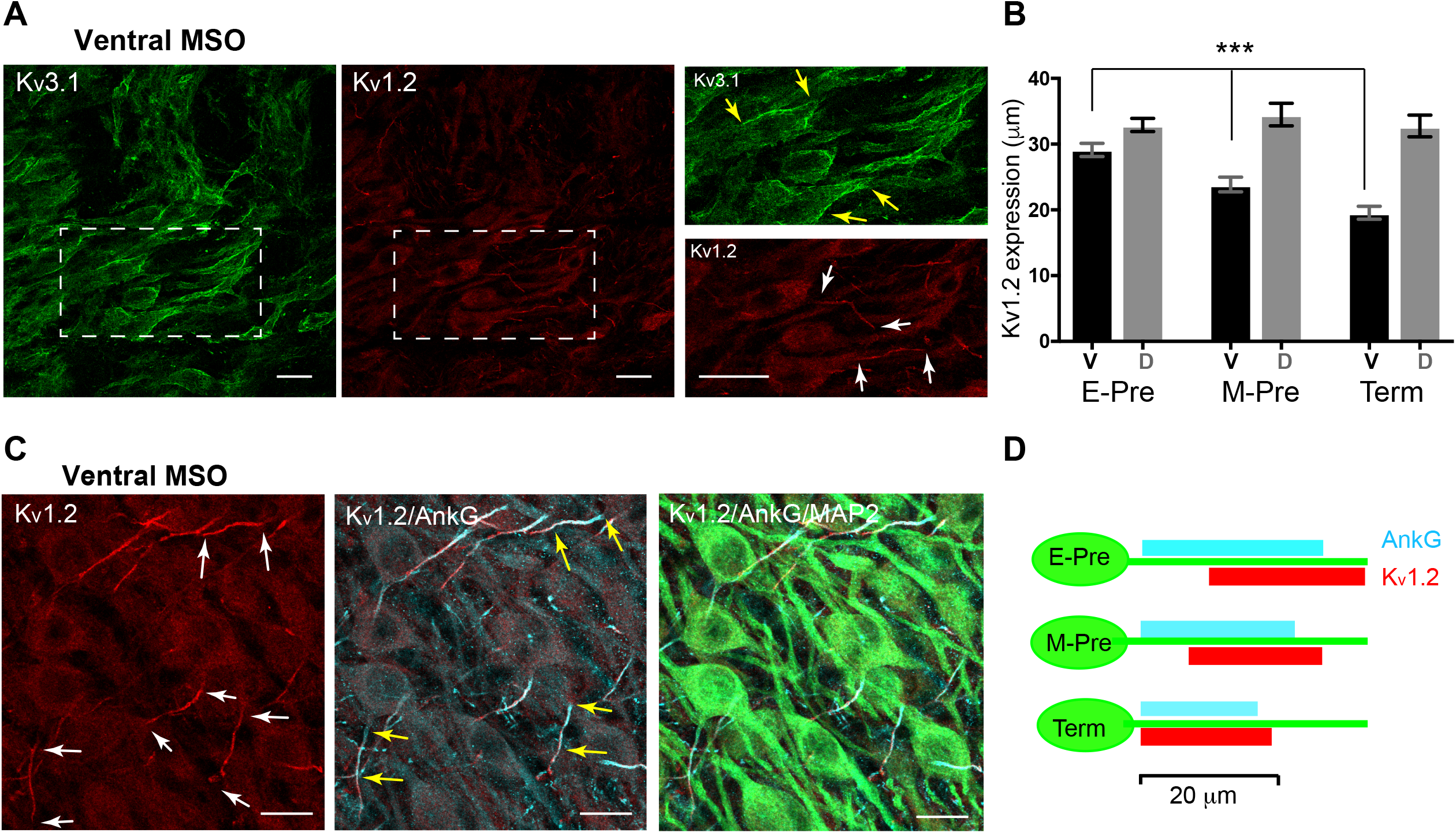
Voltage-activated K^+^ channel expression in the MSO of baboon neonates. **A.** Expression of K_v_3.1 (green) and K_v_1.2 (red) channels in the ventral MSO from M-Pre neonate. (Right) Magnified images of MSO neurons immunolabelled by K_v_3.1 and K_v_1.2 in boxed area in A. Yellow arrows indicate Kv3.1 expression on somata and dendrites of MSO neurons and white arrows indicate K_v_1.2 expression in the axon of MSO. Scale = 20 μm. **B.** Changes in the length of K_v_1.2 expression on MSO axons in ventral (V) and dorsal (D) MSO in E-Pre, M-Pre, and Term neonates. Data were shown as mean ± s.e.m. ***>0.001, one-way ANOVA test. **C.** K_v_1.2 expression (red) and colocalization with ankyrinG (AnkG, cyan) at the axon initial segment of an MSO neuron (MAP2, green) from an E-Pre baboon neonate. Scale = 20 μm. **D.** Schematic of K_v_1.2 expression pattern along the proximal axon of MSO neurons during fetal development.

### The AIS of MSO neurons undergoes dynamic changes after hearing onset in the uterus

Intrinsic excitability of MSO neurons is important for dictating neuronal response to synaptic inputs and for output signaling. Presynaptic inputs can influence the structural development of axonal domains in the avian auditory brainstem^31^. We examined structural changes in the AIS, where voltage-gated Na^+^ channels are clustered, of MSO neurons throughout gestational development. Our previous study in the mouse auditory brainstem found the structure of the AIS in MNTB neurons depends on their functional topographic location along the tonotopic axis, aligning high- to low-frequency sound-responding neurons (HF or LF neurons). Specifically, HF neurons dramatically undergo structural remodeling during early postnatal development^32^. In M-Pre and Term neonate MSO, AIS length of MSO neurons depended on the topographic location along the dorsal-ventral axis (**Figure 6A**). Ventral MSO neurons responding to high-frequency sound had a much shorter AIS compared with dorsal MSO neurons in Term baboon neonates (12.16 ± 0.876 μm, n = 28 in dorsal vs 23.06 ± 1.007 μm, n = 21 in ventral, **Figure 6B**). During gestational development, the AIS of ventral MSO neurons became significantly shorter (24.80 ± 0.422 μm, n = 61 in E-Pre, 21.02 ± 0.499 μm, n = 49 in M-Pre, and 23.06 ± 1.007 μm, n = 21 in Term, p<0.001, ANOVA), whereas AIS length of dorsal MSO neurons remained relatively stable (**Figure 6B**). In the uterus sound experience after hearing onset influences the refinement of the AIS of MSO, which is critically important to generate action potentials and contributes to output signaling of the MSO along the tonotopic axis.

**Figure 6.**
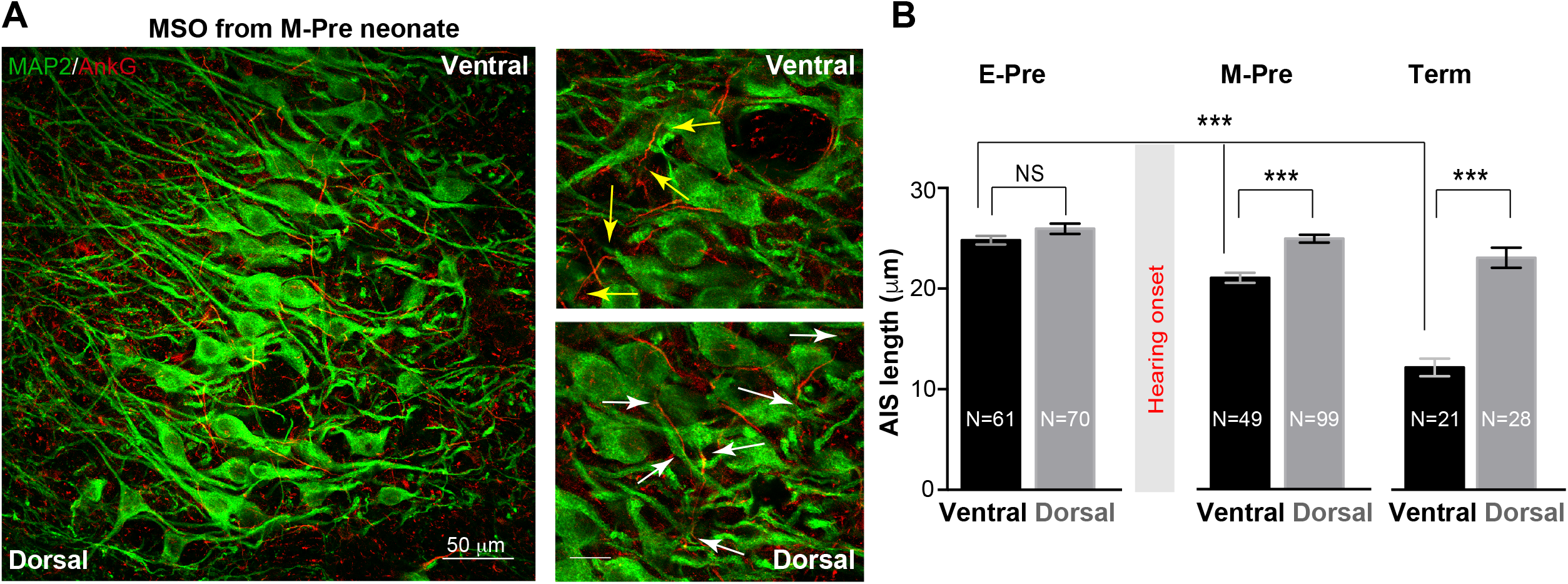
Refinement of the axon initial segment of MSO neurons at different gestational ages of baboon neonates. **A.** AnkG and MAP2 expression in ventral MSO from M-Pre neonates. The image includes 50% position (from the dorsal edge) of the MSO. Magnified images from ventral MSO and dorsal MSO immunostained with MAP2 and AnkG. Arrows indicate the length of AnkG expression. **B**. AIS length (AnkG) in the dorsal MSO and the ventral MSO at different GA. Hearing onset is between 67% (E-Pre) and 77% (M-Pre). Data were shown as mean ± s.e.m. ***>0.001, one-way ANOVA and post-hoc Tukey test.

### Effect of the NICU’s unnatural auditory environment on MSO development in preterm baboons

Most preterm newborns experience an auditory environment in the NICU, which is markedly different from the uterus^33, 34^. We investigated how altered auditory environment impacts development of morphologic features of MSO neurons in preterm baboon neonates. We compared cellular structures of MSO neurons from baboon neonates with a NICU stay (14 days after birth at 126d GA) and from age-matched baboon neonates delivered at 140d GA (M-Pre). The size of MSO (319,884 ± 18,137 μm^2^, n = 18) was not significantly different from the age-matched group at 77% GA (283,045 ± 29,823 μm^2^, n = 7). There was also no difference in soma diameter of MSO neurons, indicating that overall development of MSO was not influenced by the isolated auditory environment. However, NICU experience significantly altered the AIS length of ventral MSO neurons, thus affecting the tonotopic gradient of the AIS length. The AIS of ventral MSO neurons was much longer in neonates with a NICU stay than in the uterus (23.52 ± 0.422 μm, n=91 in the NICU vs 21.10 ± 0.508 μm, n=37 in the uterus, p=0.0013, unpaired t-test). However, there was no difference in dorsal MSO neurons (26.37 ± 0.424 μm, n=115 in the NICU vs 25.46 ± 0.432 μm, n=77 in the uterus, p=0.1511, unpaired t-test, **Figure 7A-B**). Therefore, in baboon neonates with auditory experience in the uterus, the MSO showed a clear tonotopic alignment of the AIS (shorter in ventral MSO neurons and longer in dorsal MSO neurons) at 77% GA, but neonates with a NICU stay did not. This result suggests that deprivation of maternal sound input may delay or impair the developmental refinement of the AIS of ventral MSO in the NICU, resulting in an elimination of the tonotopic segregation of the MSO.

**Figure 7.**
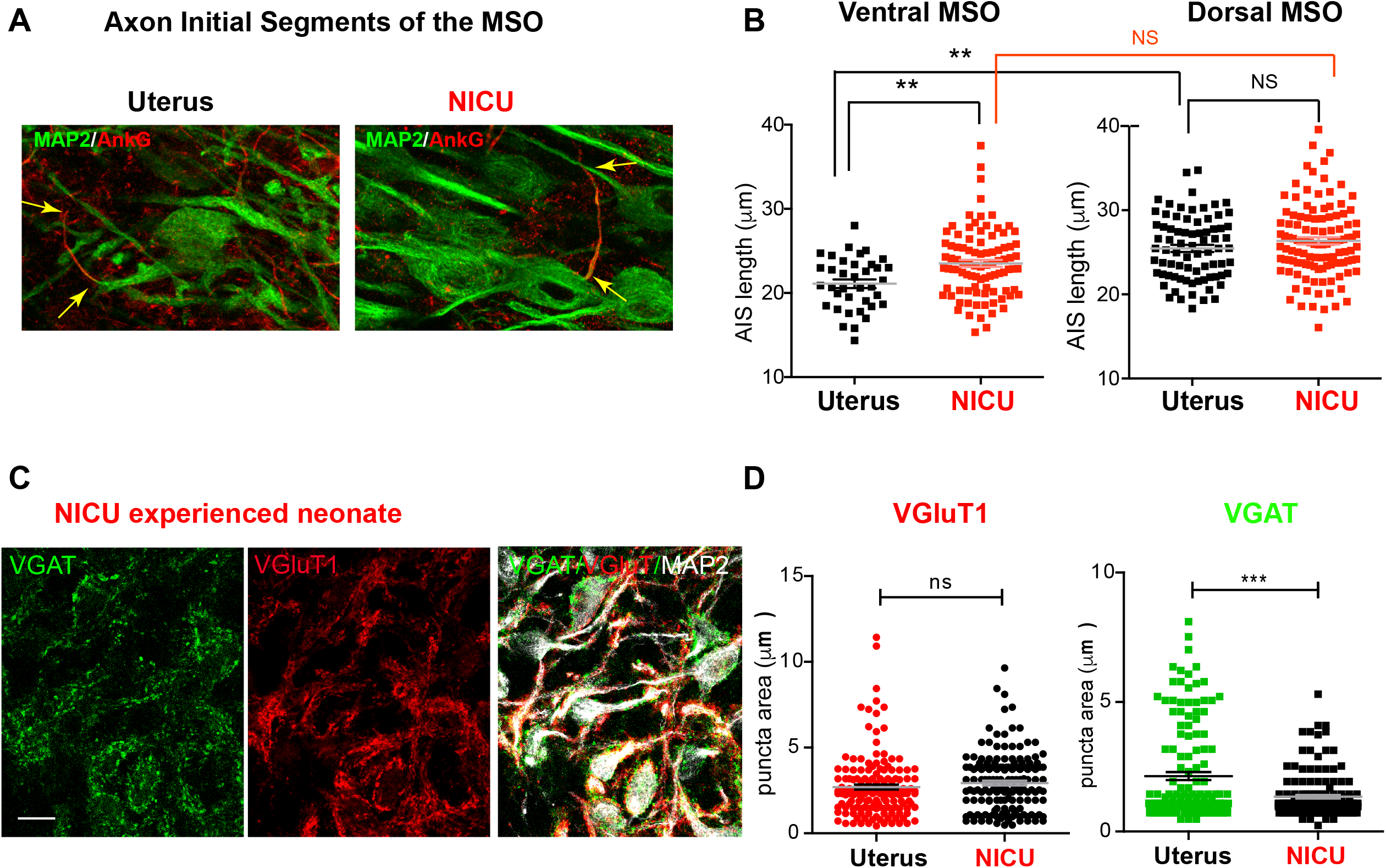
Two-week NICU stay influences structural development of the MSO in baboon neonates. **A.** Axon initial segments of MSO neurons were immunolabeled with AnkG (red) and MAP2 in (i) neonates delivered normally at 140d GA (with maternal auditory experience in uterus; Uterus), and (ii) age-matched neonates delivered at 126d with a 2-week postnatal stay in the NICU (NICU). **B.** Comparison of AIS length of MSO neurons in the ventral and dorsal MSO in the two groups (Uterus vs NICU). **C.** Immunostaining for VGluT1 (red) and VGAT (green) puncta on MSO neurons (MAP2) in neonates with a NICU stay. **D.** VGluT1 (red) and VGAT (green) puncta area in the MSO from neonates with a NICU stay (NICU) vs in the uterus (Uterus). **>0.01 and ***>0.001, unpaired t-test.

In addition to axonal structure of MSO neurons, altered sound experience after hearing onset significantly affected the development of synaptic input to the MSO. In age-matched baboon neonates, the VGAT puncta area indicating inhibitory inputs to MSO neurons was significantly reduced in baboon neonates with a NICU stay, compared with neonates in the uterus (1.34 ± 0.065 μm^2^, n= 157 in the NICU vs 2.14 ± 0.152 μm^2^, n=154 in the uterus, p=0.0088, unpaired t-test). However, there was no significant difference in VGluT1 puncta area (2.93 ± 0.136 μm^2^, n= 158 in the NICU vs 2.69 ± 0.142 μm^2^, n=159 in the uterus, p=0.0663, unpaired t-test, **Figure 7C-D**). These results indicate that a NICU stay entailing an unnatural sound environment may affect both the development of inhibitory input to the MSO and the refinement of axonal structures in MSO development.

## Discussion

The morphologic and physiologic properties of MSO neurons have well been described in rodents such as gerbils and mice ^17, 27, 29^. However, most studies using rodents have focused on postnatal development and/or the effect of auditory experience after hearing onset around postnatal day 12 (P12). Here we examined the structural properties of the MSO during fetal development in baboon neonates delivered at different gestational ages and in a NICU. Axonal and synaptic structures of MSO neurons and the overall shape of the MSO nucleus underwent profound refinement in the uterus during the last trimester after hearing onset at around 70% of GA.

MSO neurons in baboon neonates have a larger soma and more complex primary dendrites as a function of longer gestational ages. In a previous study in gerbils, one or three primary dendrites originated from the soma of MSO neurons had a number of small branches on the distal dendrites. The number of distal dendritic branches per cell decreased over age (from P8 to P15), indicating less complex dendrites ^27^. Our contradictory findings in baboon neonates could be caused by differences in species or developmental ages. In baboon neonates, MSO nuclei change from a round shape in E-pre to a thin column shape in Term neonates during the third trimester of gestation (~60 gestational days), whereas the overall shape and size of MSO nuclei in gerbils do not change based on postnatal ages ^27^. This could reflect different fetal and postnatal developmental effects on the MSO in different species. In addition, our complexity analysis of MSO neurons focused on their somata and primary dendrites, and excluded dendritic branches. In E-Pre neonates, most MSO neurons had distinct bipolar structures with two primary dendrites extending in a medial-lateral orientation. In Term neonates, MSO neurons showed a smaller soma and more primary dendrites stretching in multiple directions. These conflicts between gerbil and baboon neonates could be due to differences in the fetal and postnatal development of the MSO in different species or due to differences in relative developmental ages studied. It should be noted that in this study, morphologic analysis was based on MAP2 immunolabelling for multiple neurons instead of post-recording immunostaining for a single cell. Thus, it is possible to underestimate sizes of the distal structures of MSO dendrites located far away from the stomata in baboon neonates.

After hearing onset, spatial refinement of excitatory and inhibitory synaptic inputs on the dendritic and somatic surface is critical for MSO neurons to allow synaptic integration for fine-tuning of the coincidence mechanisms ^15, 16^. Similar to the MSO in gerbils ^15^, the distribution of inhibitory inputs in MSO was refined after hearing onset in baboons, in the uterus and throughout fetal development ^25^. Before hearing onset in E-pre neonates, both VGluT1 and VGAT signals were detected at the somata and dendrites without a distinguishable pattern. After hearing onset, VGAT signals, indicating inhibitory inputs, were concentrated on the alignment of MSO somata throughout Term neonates, whereas VGluT1 signals were concentrated on dendrites. Our results in baboon neonates support the concept that auditory experience in the uterus promotes spatial refinement of excitatory and inhibitory inputs on MSO neurons, allowing them to detect interaural time differences in low-frequency hearing mammals ^15^. In addition to the spatial distribution of synaptic inputs, the VGluT1 and VGAT puncta on MSO neurons significantly enlarged after hearing onset during the last trimester in the pregnancy. Our ultrastructural analysis using electron microscopy supports the idea that the increased size of VGluT1 and VGAT puncta may be due to increased synaptic vesicle numbers around the active zone in the individual boutons. Although inhibitory and excitatory synapses were not distinguishable in these analyses, most synapses had a similar staining patterns during fetal development.

MSO neurons play a role in detecting coincidences within interaural time differences in the range of a few tens of microseconds for computing sound location. To achieve this task with remarkable acuity, voltage-activated K^+^ (K_v_) channels critically contribute to setting a short integration time constant of the membrane and a typical spiking pattern ^17, 27, 29, 30^. In baboon neonates, a single spike of MSO neurons in response to a long-depolarizing current injection displayed a relatively high threshold and a lower amplitude, similar to findings in rodents. In gerbils and mice, the amplitude of spikes and input resistance in MSO neurons progressively decreased after hearing onset ^17, 30^, whereas we observed no significant difference in baboons between E-Pre and M-Pre neonates. In gerbils, low voltage-activated potassium (K_v_1) channels and high voltage-activated (K_v_3) channels were detected along the MSO neuron from soma to dendrites ^29^. In mice, K_v_1 currents determine the AP threshold and firing pattern and K_v_3 currents contribute to AP width ^17^. In baboons, we detected K_v_3.1 channel along the MSO neurons from the soma to the primary dendrites, whereas K_v_1.2 channels appear to be restricted to the axon initial segment of MSO neurons. The lengths of K_v_1.2 expression progressively reduced along the axon initial segment of MSO neurons during fetal development. In rodents, K_v_1 currents were increased during postnatal development; this could drive developmental changes in intrinsic properties of MSO neurons, resulting in a lower amplitude and a higher threshold of APs ^17, 30^. We observed no significant changes in AP waveforms in MSO neurons between E-Pre and M-Pre neonates (data not shown). Future studies will be required to elucidate whether a shorter length of K_v_1.2 expression in baboon MSO is directly associated with reduced K_v_1 currents and/or changes in AP shape during fetal development including Term neonates.

Preterm infants admitted to the NICU frequently show some abnormalities in auditory brainstem responses ^35^. The transduction of sound information along the auditory pathway affects activity-dependent circuitry formation during a critical period for auditory development, i.e., when the nervous system is especially sensitive to certain environmental sound stimuli ^13^. Preterm newborns experience a drastic change from the maternal environment of the uterus to the NICU. The missing of maternal sound inputs (e.g. heart-beating sound in the uterus) in the NICU coincided with a significantly delayed or impaired synaptic and axonal development of MSO neurons that normally occurs during the critical period in baboon neonates. Our findings support that the isolated and unnatural auditory environment in the NICU critically impacts the structural refinement of neurons and synapses in the MSO that occurs during perinatal development. The NICU generally has a noisy environment with relatively high-frequency sound (>500 Hz broad spectrum and 70-75 dB in daytime), which derived from routine hospital noise levels ^34^. Thus, future studies need to evaluate how either the noisy environment in the NICU or maternal sound deprivation separately or synergistically contributes to the anatomic and functional development of the MSO in baboon neonates.

## Materials and Methods

### Animals

We used 15 baboon brains, including 3 term and 8 preterm baboons (either sex), and 4 preterm baboons (delivered at 126 days of gestational ages and kept in the NICU for postnatal 14 days) obtained from the laboratory of Dr. Cynthia L. Blanco (Department of Pediatrics, University of Texas Health Science Center San Antonio [UTHSCSA]). Baboons were used in accordance with approved UTHSCSA Institutional Animal Care and Use Committee protocols. Preterm baboons were delivered at the Texas Biomedical Research Institute in San Antonio. Preterm baboon care has been described in detail (Blanco et al., 2013). Briefly, term baboons were delivered naturally (at 180 d GA), whereas preterm baboons were delivered via cesarean section at 125 ± 2 d GA (67% of GA) and at 140 ± 2 d GA (77% of GA). Preterm baboon neonates were intubated immediately after birth and chronically ventilated for a planned survival of 14 d in the NICU (with sound attenuation, < 40 dB, in isolated room). Central intravenous lines were placed shortly after birth for fluid management and parenteral nutrition. In the NICU, preterm baboons were treated with corticosteroids, antibiotics, ketamine, vitamin K, and blood transfusions (Blanco et al., 2013). Any possible effects of extreme preterm birth independent from NICU experience were not assessed in this study, as NICU interventions were necessary for the survival of the neonates.

### Slice preparation

Brainstem slices were prepared from term and preterm baboon brainstems after necropsy. Brainstems were immersed in ice-cold low-calcium artificial CSF (aCSF) containing the following (in mM): 125 NaCl, 2.5 KCl, 3 MgCl_2_, 0.1 CaCl_2_, 25 glucose, 25 NaHCO3, 1.25 NaH_2_PO_4_, 0.4 ascorbic acid, 3 myo-inositol, and 2 Na-pyruvate, bubbled with carbogen (95% O_2_, 5% CO_2_; pH=7.3-7.4; osmolality of 310-320 mOsm). Transverse brainstem slices (200 μm thick) were prepared using a microtome (VT1200S, Leica). After cutting, the slices were transferred to an incubation chamber containing normal aCSF bubbled with carbogen and maintained at 35°C for 30 min and thereafter at room temperature. Normal aCSF was the same as the slicing aCSF, but with 1 mM MgCl_2_ and 2 mM CaCl_2_.

### Electrophysiology

Whole-cell patch-clamp recordings were performed in normal aCSF at room temperature (22-24°C). The pipette solution contained (in mM): 130 K-gluconate, 20 KCl, 5 Na_2_-phosphocreatine, 10 HEPES, 5 EGTA, 4 Mg-ATP, and 0.3 GTP, pH adjusted to 7.3 with KOH. EPSCs or action potentials (APs) were recorded in normal aCSF using the voltage or current-clamp mode of the EPC-10 (HEKA Electronik, Lambrecht/Pfalz, Germany). Patch electrodes had resistances of 2.5-3 MΩ and the initial uncompensated series resistance (Rs) was < 20 MΩ. Recordings were not corrected for the predicted liquid junction potential of 11 mV. APs were elicited by current injection in current-clamp mode. Data were filtered at 2.9 kHz and digitized at a 10-50 μs sampling intervals.

### Immunohistochemistry

After electrophysiologic recordings, brainstem slices (150-200 μm) were fixed with 4% (wt/vol) paraformaldehyde in phosphate buffer saline (PBS) solution for 30 minutes. Free-floating sections were blocked in 4% goat serum and 0.3% Triton X-100 in PBS for 1 hour. Primary antibodies were guinea pig polyclonal anti-vesicular glutamate transporter 1 (VGluT1, 1:3000, Chemicon), anti-vesicular GABA transporter (VGAT, 1:1000, Synaptic Systems Ca# 131-002), anti-microtubule associated protein 2 (MAP2, 1:200, Millipore), mouse monoclonal anti-ankyrinG (AnkG; 1:100; UC Davis/NIH NeuroMab Facility Cat# 75-146, RRID:AB_10673030), mouse monoclonal anti-K_v_1.2 (1:250; UC Davis/NIH NeuroMab Facility Cat# 75-008, RRID:AB_2296313), mouse monoclonal anti-K_v_3.1 (1:250; UC Davis/NIH NeuroMab Facility Cat# 73-041). After 3 washes with PBS containing 0.1% Tween 20, slices were incubated with different Alexa-488 goat anti-mouse IgG1 or 568 goat anti-mouse IgG2b or 568 goat anti-rabbit or 647 goat anti-guinea pig secondary antibodies (1:1000; Invitrogen) accordingly for 2 hours at room temperature. Slices were then rinsed with PBS containing 0.1% Tween 20 and were cover-slipped using mounting medium (Vectashield; Vector Laboratories). Stained slices were viewed on a confocal laser-scanning microscope (Carl Zeiss LSM-710) at 488, 568, and 633 nm using a pinhole of 1 AU and 20x (0.8 NA), 40x (oil-immersion, 1.30 NA) objectives. Z-stack images were acquired at a digital size of 1024 x 1024 pixels with optical section separation (*z* interval) of 0.5 um. Images were imported into Fiji (Image J) ^36^, ZEN (Carl Zeiss), and CellProfiler for analyses. In VGluT1 and VGAT puncta, regions of interest were selected based on MAP2 staining and overlapped with Vglut1 and VGAT staining using CellProfiler. The numbers and sizes of puncta were analyzed using the analysis modules “IdentifyPrimaryObjects and IdentifySecondaryObjects” with settings of 1) typical diameter of objects in pixel units (Min., Max.) = 3, 10; 2) method to distinguish clumped objects = intensity; 3) discard objects outside the diameter range = yes; 4) discard objects touching the border of the image = yes; and 5) threshold=0.2~0.3. For AIS, 2D compressed Z-stack images were analyzed using the Fiji program as we have described previously ^32^. Briefly, the AIS was determined as the proximal axon where the fluorescence intensity of ankyrin G staining was >10% of the peak signal. Length of the AIS was measured by the segmental line profile in the Fiji program.

### Theoretical fractal dimension analysis

The complexity of dendritic arbors was examined by determining fractal dimensions ^26^. The box-counting method was applied for fractal analysis using fractal3e software (http://cse.naro.affrc.go.jp/sasaki/fractal/fractal-e.html). The number (N) of squares (boxes) in a square grid contacted by the dendrites was counted, as grids with decreasing sizes of squares are placed over the cell. The fractal dimension is depicted as the relationship between increasing N and decreasing size of squares, with larger numbers indicating increased complexity ^27^.

### Transmission electron microscopy

Brains were removed and 400-μm-thick samples of brainstem MSO area were dissected out followed by primary fixation in 1% glutaraldehyde/4% paraformaldehyde. Further processing was performed by the UTHSCSA Electron Microscopy Lab. Briefly, each brainstem was post-fixed with 1% Zetterqvist’s buffered osmium tetroxide, dehydrated, and embedded in PolyBed resin at 80°C in an oven. Tissue containing the MSO was cut into 90-nm ultrathin sections and placed on copper grids. The sections were then stained with uranyl acetate and Reynold’s lead citrate. The samples were imaged on a JEOL 1400 electron microscope using Advanced Microscopy Techniques software. A total of 20-40 synapses was analyzed from 3 baboons for each age group. The number of docked vesicles per the active zone was measured for each synapse at a final magnification of 80,000×. The active zone was defined as the dark presynaptic density contacting the postsynaptic density. Docked and clustered vesicles were defined as those within 10 nm and 200 nm of the presynaptic active zone, respectively ^37^.

### Analysis and statistics

Electrophysiologic data were analyzed and presented using Igor Pro (Wavemetrics, OR). All statistical analyses were performed in Prism (GraphPad Software). Normality of datasets was analyzed using the D’Agostino and Pearson’s omnibus test. Parametric or non-parametric tests were carried out accordingly. α values were set to 0.05, and all comparisons were two-tailed. To compare two groups, unpaired t-tests or Mann-Whitney U tests were carried out. For three or more groups, one-way ANOVA with post-hoc Tukey’s multiple comparison test was used. Significance was set at *P* <0.05. Data were shown as the mean ± standard error of the mean (s.e.m.) with *n* values representing the number of animals per experimental group or the number of neurons per group where indicated. In box and whisker plot, boxes indicate 25-75% interquartile range and horizontal lines in boxes indicate median. Whiskers show 5% ~ 95% range and dots show outliers that reside outside the whisker range. Differences were deemed statistically significant when *p* values were <0.05.

## Acknowledgements

This work was supported by a grant from the National Institute on Deafness and Other Communication Disorders (NIDCD, R01 DC03157), a Southwest National Primate Research Center (SNPRC) Pilot Grant, and a Children’s Health Pilot Program grant (UTHSCSA), all to J. H. Kim.

## Acknowledgements

This work was supported by a grant from the South National Primate Research Center (SNPRC) Pilot Grant and Children’s Health Pilot Program (UTHSCSA) to J. H. Kim.

